# Deposition of microplastics associated with bioaccumulation of heavy metals in human lungs of smokers: Implications of adsorption and mobilization of metals via microplastics

**DOI:** 10.64898/2025.11.30.691449

**Authors:** Gagandeep Kaur, Faizan Imam, Felix Effah, Aerlin D Decker, Marcus A Garcia, Matthew J Campen, Irfan Rahman

**Author notes:** Correspondence should be addressed to: Irfan Rahman, Ph.D., Department of Environmental Medicine University of Rochester Medical Center Box 850, 601 Elmwood Avenue Rochester 14642, NY, USA.

## Abstract

**Background:** Airborne micro- and nano plastics (MNPs), generated from synthetic textiles, vehicle tires and plastic wastes, are omnipresent in the environment and known to deposit in human lungs. Bioaccumulation of MNPs in lung tissues has been studied, but effect of tobacco smoking on deposition is not well understood. Tobacco smoke is known to contain heavy metals which can get adsorbed to the MNP surface, leading to increased environmental bioavailability. We hypothesized that smoking leads to enhanced chelation of heavy metals with airborne MNPs leading to accentuated lung bioaccumulation, thereby leading to lung pathologies.

**Methods:** Age (20-75 years) and sex-matched archival lung tissue samples from smokers (n = 9) and non-smokers (n = 10) were procured and quantified to assess: (a) the levels of MNPs by pyro-GC/MS and (b) heavy metals by ICP-MS. Furthermore, to establish if there exists a correlation between MNP deposition and disease occurrence in current smokers, tissue samples from age-matched patients with COPD (n = 4) and IPF (n = 4) underwent pyro-GC/MS analyses.

**Results:** A relatively higher level of MNPs were found in the lungs of smokers (43.67 mcg/gm) as compared to non-smoker (34.53 mcg/gm) upon pyro-GC/MS analyses. We observed a marked increase in the levels of polyethylene (PE; p< 0.0076), polycarbonate (PC; p< 0.0947) and nylon6 (N6; p < 0.0653) polymers in the lungs of smokers as compared to non-smokers. Metal analyses showed a significant (p <0.0001) increase in the levels of cadmium (Cd) in the lungs of smokers with the levels of deposited Cd being more pronounced for female smokers. Importantly, we observed a strong correlation (p < 0.05) between the amounts of accumulated PE and levels of Cd in the lungs. Pyro-GC/MS analyses did not reveal any relation between MNP deposition and disease occurrence in our study. Furthermore, little to no change in the level of deposited MNPs was observed in the lungs of current versus ex-smokers in disease cohorts.

**Conclusions:** Overall, our results show that smoking promotes bioaccumulation of certain MNPs (PE) with heavy metals (Cd) which may lead to adverse lung effects in smokers.

## Introduction

Micro- and Nano plastics (MNPs) comprising particles from the micron to submicron range are omnipresent in the modern world; detected in food products, drinking water and air (Xu, Lin et al. 2022, Vasse and Melgert 2024). Reports show the presence of MNPs in the food web (Mamun, Prasetya et al. 2023) and bioaccumulation in human tissues, including liver (Horvatits, Tamminga et al. 2022), colon (Ibrahim, Tuan Anuar et al. 2021), kidneys (Massardo, Verzola et al. 2024), lungs (Amato-Lourenço, Carvalho-Oliveira et al. 2021, Jenner, Rotchell et al. 2022, Lu, Li et al. 2023) and even brain (Nihart, Garcia et al. 2025) and placentae (Ragusa, Svelato et al. 2021, Garcia, Liu et al. 2024, Jochum, Garcia et al. 2025). However, research into the implications of bioaccumulating MNPs on health and disease is not well understood.

Since its first citation as a major source of pollution in 2004 (Thompson, Olsen et al. 2004), much effort has been put into characterizing and exploring the effects of MNPs in water and food sources (Jiang, Kauffman et al. 2020, Xu, Lin et al. 2022, Habumugisha, Zhang et al. 2024, Vasse and Melgert 2024). But the harms associated with other routes of exposure, especially air-borne, have gained interest recently (Vasse and Melgert 2024). A 2022 report by the World Health Organization has approximated a daily intake of 3000 microplastic particles upon inhalation of 15 m^3^ air (WHO 2022). However, it is important to acknowledge that the average human exposures vary depending upon the location on earth (more/less polluted; rural/urban), site (indoor/outdoor), size, shape and aerodynamic diameter of the particles present in the air at the time of measurement (Dris, Gasperi et al. 2017, Liu, Wang et al. 2019, WHO 2022, Vasse and Melgert 2024). Particles with an aerodynamic diameter between 2.5 to 100 μm are considered inhalable, with smaller sized (< 2.5 μm) particles having capabilities to penetrate deeper into the peripheral lung regions (Donaldson and Tran 2002, Vasse and Melgert 2024). The common sources of airborne MNPs include synthetic textiles (Periyasamy and Tehrani-Bagha 2022, Akyildiz, Fiore et al. 2024), road wear (Järlskog, Jaramillo-Vogel et al. 2022, Yoo, Park et al. 2025), vehicle tires (Saladin, Boies and Giorio 2024, Yoo, Park et al. 2025), incineration of waste (Mokammel, Naddafi et al. 2025, Su, Wang et al. 2025), and burn pit smoke (Kim, Warren et al. 2021, Penuelas and Lo 2024). Characterizing the chemical composition of the inhaled MNPs found in the peripheral lung tissues reported the predominance of polypropylene (PP) and polyethylene (PE) and polyethylene terephthalate (PET) polymers in humans (Amato-Lourenço, Carvalho-Oliveira et al. 2021, Jenner, Rotchell et al. 2022, Wang, Lu et al. 2023). In fact, a 2021 study characterizing the lung toxicity and chemistry of burn pit smoke emissions identified plastic waste to be the most significant contributor of particulate matter that resulted in the production of the highest sum of toxic volatile organic compounds (VOCs) upon combustion (Kim, Warren et al. 2021).

Studies suggest the toxic effects of microplastics, providing evidence of oxidative stress and DNA damage *in vitro* (Zhu, Jia et al. 2020, Milillo, Aruffo et al. 2024, Wei, Chen et al. 2025), immune-modulation and metabolic imbalance *in vivo* (Woo, Seo et al. 2023, Garcia, Romero et al. 2024, Wang, He et al. 2025) and lack of function in organoid models (Winkler, Cherubini et al. 2022). However, the scope of these studies is quite limited. Most of the current studies have used polystyrene (PS)-microspheres to test the MNP toxicity (Dong, Chen et al. 2020, Florance, Chandrasekaran et al. 2022, Merkley, Moss et al. 2022, Garcia, Romero et al. 2024, Milillo, Aruffo et al. 2024, Wang, He et al. 2025). However, PS does not form the major type of microplastic contaminating the ingestible or inhalable fraction of MNPs (Li, Tao et al. 2023, Wright, Cassee et al. 2024, Ragu Prasath, Sudhakar and Selvam 2025). Furthermore, the inhaled MNPs exhibit a variety of shapes, including fibers (long and thin), fragments (due to the weathering process), foams, and films (El Hayek, Castillo et al. 2023, Ward, Gordon et al. 2024, Wright, Cassee et al. 2024). Most of the current studies suggesting the harmful effects of inhaled MNPs *in vitro* have been conducted using A549 cells (Zhu, Jia et al. 2020, El Hayek, Castillo et al. 2023, Woo, Seo et al. 2023, Milillo, Aruffo et al. 2024, Michelini, Mawas et al. 2025, Wang, He et al. 2025). Though this model is well characterized for toxicity studies (Foster, Oster et al. 1998), it does not represent the alveolar type II cells due to its origin from adenocarcinoma (Swain, Kemp et al. 2010). Proof exists supporting the development of lung diseases, including asthma (Hussey 1976, Gehring, Wijga et al. 2020), pneumoconiosis (Mastrangelo, Saia et al. 1981), interstitial lung diseases (ILD) (Lilis 1980) and lung cancer (Girardi, Barbiero et al. 2022, Justeau, Gervès-Pinquié et al. 2022). Still, direct evidence showing mechanistic toxicities of MNPs resulting in physiological effects is yet to be explored.

Evidence from aquatic environments has suggested that microplastics act as strong magnets for heavy metals (Liu, Shi et al. 2021). Interaction between microplastics and heavy metals increases the risk of retention of these conjugates in the environment. It raises the possibility of them being more bioavailable to enter the food chain (Akhbarizadeh, Moore and Keshavarzi 2018, Abidli, Akkari et al. 2021). We hypothesized that metal-MNPs chelated forms can be easily transported to different organs via biological fluids. It is thus expected that the combined toxic effects of microplastics and metals can be more severe than the effects of either pollutant alone, potentially causing greater harm to pulmonary epithelium and extracellular matrix via induction of oxidative stress and immune-inflammatory responses. In this regard, heavy metals like cadmium (Cd), arsenic (As) and lead (Pb) are of major concern as these are abundant in tobacco smoke (Piadé, Jaccard et al. 2015, Matt, Quintana et al. 2021) and are known to accumulate in lung tissues from smokers (Chen, Peng et al. 2016, Pinto, Cruz et al. 2017).

Considering this, we quantified and characterized the major MNPs and heavy metals in the lungs of healthy smokers and non-smokers in this study. We also studied the correlation between the bioaccumulation of MNPs and toxic heavy metals within the human lungs, to establish a potential link between bioaccumulation of metal-MNP chelates and the occurrence of various pathologies in the lung.

## Materials and Methods

### Ethics statement

The experiments were carried out according to the standards and guidelines approved by the University of Rochester Institutional Biosafety Committee. Office of Human Subject Protection (RSRB) at the University of Rochester Medical Center (URMC) (Study ID: STUDY00006571 dated October 5, 2021) approved the study protocols for research pertaining to human subjects. All the experiments were conducted in a blinded fashion and care was taken to employ a robust and unbiased approach to uphold the NIH standards of rigor and reproducibility.

### Human Sample Procurement

Human lung tissue samples from healthy non-smokers (NS) and smokers (n = 9-10 / group) were procured from two sources – (a) United Network for Organ Sharing, facilitated by the International Institute for the Advancement of Medicine (IIAM) and (b) National Disease Research Interchange (NDRI)-for this study. These are non-profit organizations that provide non-transplantable, fresh or frozen human organs/tissues for medical research. Healthy human lungs with no history of infection or chronic lung ailment were included in this study. Age (20-75 yrs)- and sex-matched samples from healthy NS and smokers were chosen and used for further downstream analyses. The clinical characteristics of the samples included in this study are presented in **Table 1**.

**Table 1:**
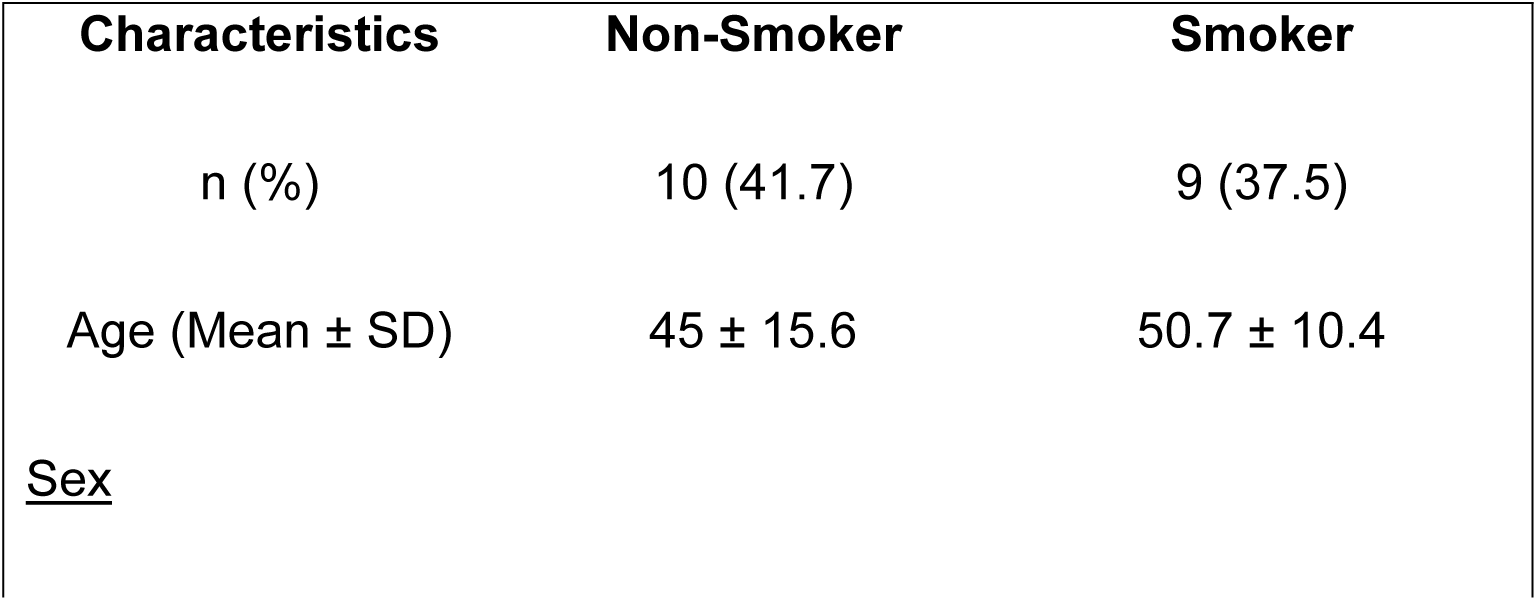
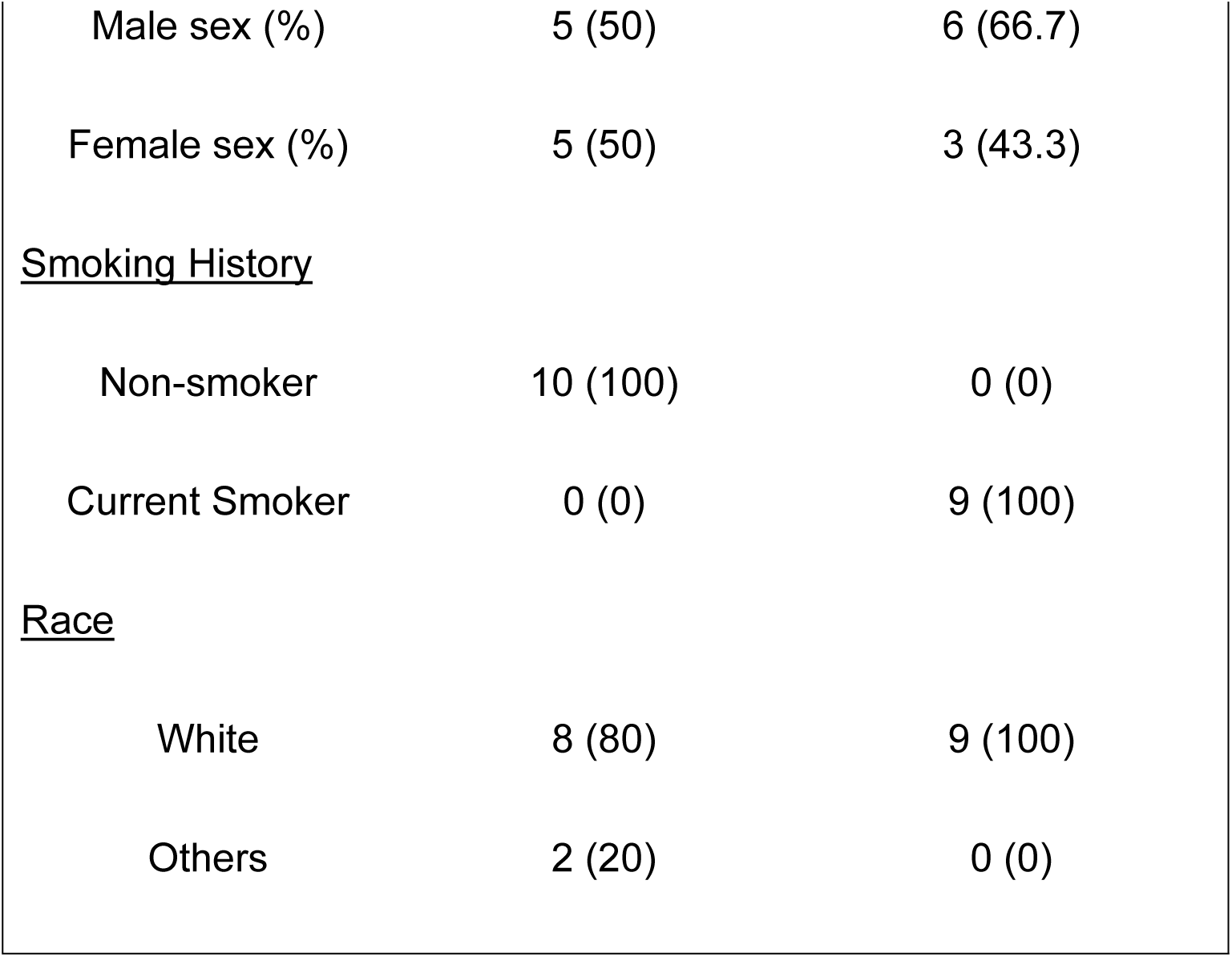
Clinical Characteristics of lung tissue samples used for microplastic.

### Microplastic Isolation

Biological samples were digested using the saponification method as described previously (Nihart, Garcia et al. 2025). In brief, approximately 400 mg of lung tissue samples were chopped into smaller chunks and transferred to glass vials for saponification using 10% potassium hydroxide (KOH) at 40°C for 72 hours with intermittent manual mixing. Following consistent and thorough digestion of the tissue chunks with no visible debris, 200 µl of 100% ethanol was added and the microplastic fraction was separated from the tissue digest by ultracentrifugation at 100,000 × g for 4 hours using an Optima Max-XP ultracentrifuge (Beckman Coulter, Brea, CA, USA). The microplastic fraction could be clearly seen as a pellet at the bottom of the tube following ultracentrifugation. The supernatant was discarded, and the pellet was washed twice with 100% ethanol to remove residual contaminants. After the ethanol washes, the pellet was transferred to cyclohexane and resuspended at 60°C for 24 hours to further eliminate residual lipids. The cyclohexane mixture was then passed through a 50 nm pore quartz filter. The filters were then air-dried overnight under sterile conditions and stored for further analyses **(Figure 1)**.

**Figure 1:**
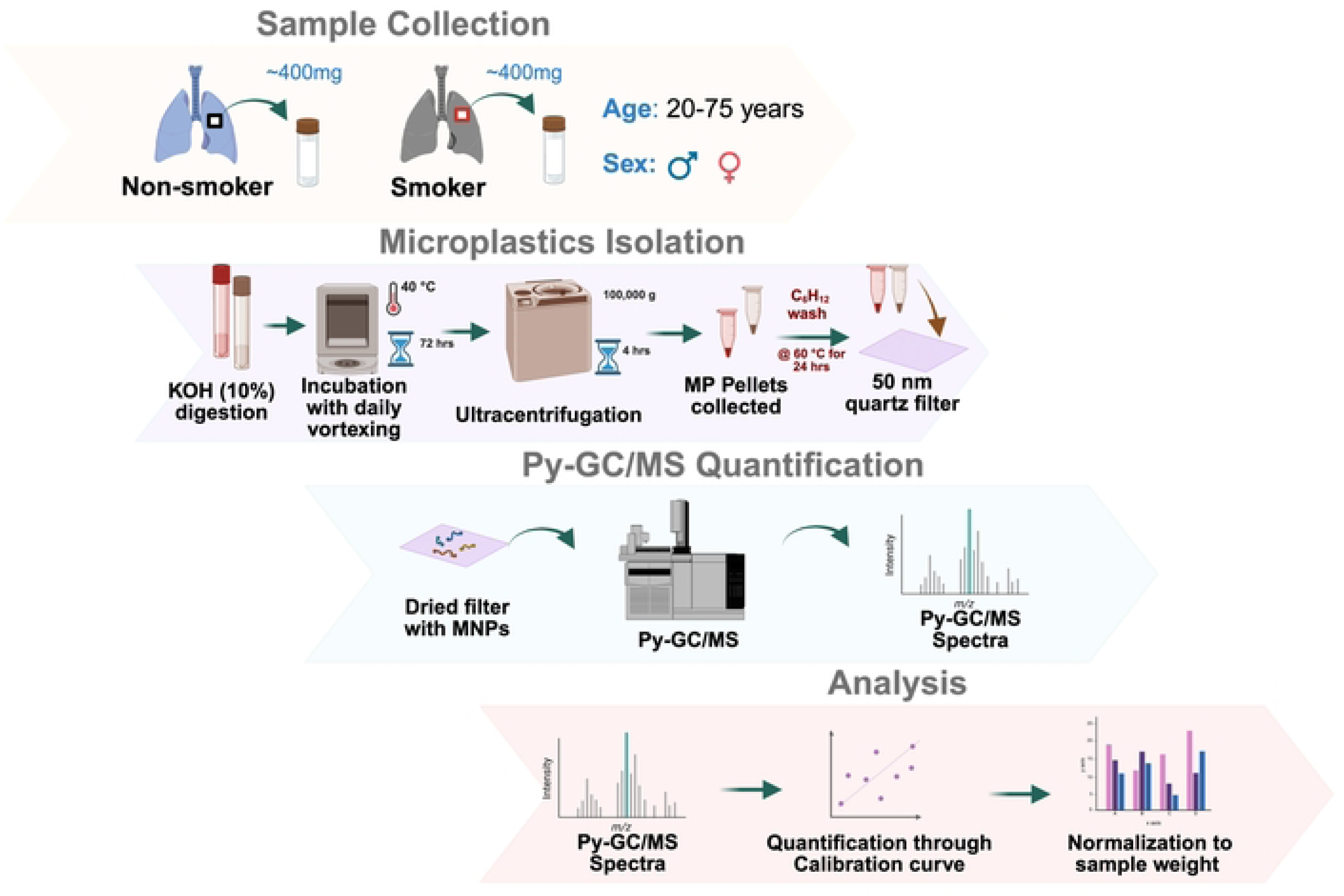
Schematics showing the method used for quantification of microplastics from lung tissues. Approximately 400 mg of lung tissue samples were collected from age (20-75 yrs) and sex-matched healthy non-smokers (NS) and smokers (n = 9-10/ group). The collected tissue samples were digested by adding 10% of potassium hydroxide (KOH) and incubated with daily vortexing at 40℃ for 72 hrs. Thereafter, the samples were ultracentrifuged at 100,000 g for 4 hrs, processed and the collected on quartz filters prior to quantification using pyrolysis-gas chromatography/mass spectrometry (Py-GC/MS). The quantified MPs were normalized using per gram of lung tissue used for digestion and plotted. The image is generated using Biorender.

### Microplastic Quantitation and Analysis

We used pyrolysis gas chromatography /mass spectrometry for quantification of microplastics using the method and instrument settings described by Nihart *et al*. (2025) (Nihart, Garcia et al. 2025). Py-GC/MS is a highly sensitive analytical technique used to identify and quantify the components of a microplastic sample by combusting solid polymers into smaller ion fragments, separating each fragment by gas chromatography, and quantifying using mass spectrometry (Santos, Insa et al. 2023). Briefly, the filter from each sample was quantitated by Py-GC/MS and compared to a microplastics-CaCO3 standard (5-point curve) containing 12 specific polymers: Polyethylene (PE), Polyvinyl chloride (PVC), Nylon 66 (N66), Styrene-butadiene (SBR), Acrylonitrile Butadiene Styrene (ABS), Polyethylene terephthalate (PET), Nylon 6 (N6), Poly(methyl methacrylate) (PMMA), Polyurethane (PU), Polycarbonate (PC), Polypropylene (PP), Polystyrene (PS). Polymer spectra were identified via the F-Search MPs v2.1 software (Frontier Labs) used for identifying the polymer spectra, and data were normalized to the weight of lung tissue used for microplastic isolation (mcg/gm) **(Figure 1)**.

### Elemental Metal Quantitation

To assess the levels of select elements in lung tissue samples from smokers’ and non-smokers’ lungs, we employed inductively coupled plasma mass spectrometry (ICP-MS) (Planeta, Kubala-Kukus et al. 2021). Approximately 80-100 mg of lung tissue samples were sent to the elemental analyses core at URMC and levels of sulphur (S), lithium (Li), beryllium (Be), sodium (Na), magnesium (Mg), aluminum (Al), silicon (Si), phosphorus (P), potassium (K), calcium (Ca), titanium (Ti), vanadium (V), chromium (Cr), manganese (Mn), iron (Fe), cobalt (Co), nickel (Ni), copper (Cu), zinc (Zn), gallium (Ga), germanium (Ge), arsenic (As), selenium (Se), rubidium (Rb), strontium (Sr), zirconium (Zr), niobium (Nb), molybdenum (Mo), ruthenium (Ru), rhodium (Rh), palladium (Pd), silver (Ag), cadmium (Cd), indium (In), tin (Sn), antimony (Sb), tellurium (Te), cesium (Cs), barium (Ba), hafnium (Hf), tantalum (Ta), tungsten (W), rhenium (Re), iridium (Ir), platinum (Pt), gold (Au), mercury (Hg), thallium (Tl), lead (Pb), bismuth (Bi), uranium (U), gadolinium (Gd), and terbium (Tb) were quantified. The levels of elemental metals were normalized to the amount (mg) of lung tissue used for quantification and plotted.

### Statistical Analysis

GraphPad Prism software, version 10.5.0, was used for plotting the graphs and performing the statistical analyses. The significance level was set to be less than or equal to 0.05. Pairwise comparisons were performed using a two-tailed Mann-Whitney U-test, and multiple comparisons using sex and smoking status as independent variable were performed using Two-way ANOVA (analysis of variance) with post hoc Tukey’s HSD (honestly significant difference) test using sex and smoking status as independent variables. Pearson’s correlation was used to establish the correlation between the levels of cadmium and microplastics (PE) identified in the matched samples.

## Results

### The average total accumulation of microplastics was higher in smokers’ lungs

This study was designed to study whether microplastic accumulation within the lungs is dependent on the smoking status of the donor. To study this, frozen lung samples from healthy sex-matched adults (20-72 years) with and without a history of smoking were procured and used for isolation and quantification of microplastics using a standardized protocol, as shown in **Figure 1**.

All 12 categories of microplastics being studied were detected within detectable limits in the lungs of both non-smokers (NS) and smokers. Polyethylene (PE), Polyvinyl chloride (PVC), Polyethylene terephthalate (PET) and Polyurethane (PU) were the most abundant categories of microplastics detected in the human lungs, on average **(Figures 2A & 2C)**. Importantly, the average total accumulation of microplastics within the lungs of smokers (43.67 mcg/gm of lung tissue) was found to be higher than that of a healthy non-smoker (34.53 mcg/gm of lung tissue) **(Figures 2B & 2D)**. **Supplementary Figure** 1 shows the distribution of the 12 MNPs tested in this study in each sample plotted with respect to the limit of detection and quantification of the technique.

**Figure 2:**
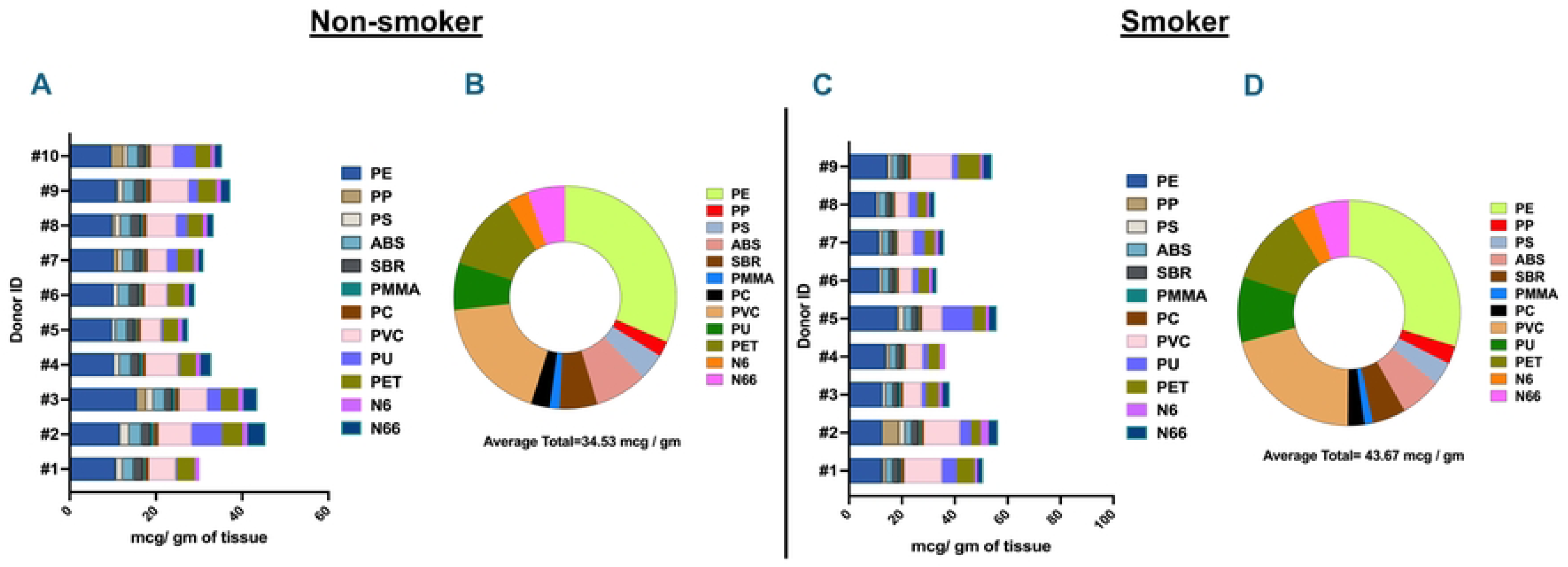
The average total of the deposited microplastics was higher in the lung tissue from smokers. Stacked Bar chart **(A&C)** and donut plot **(B&D)** showing the distribution of individual microplastics in each donor lung and the average total of each microplastic in the lungs of healthy non-smokers **(A&B)** and smokers **(C&D)**.

### Smoker’s lungs have substantial accumulation of polyethylene (PE), polycarbonate (PC) and nylon6 (N6) polymers

Upon comparing the levels of individual microplastic polymers found within the lungs of smokers and non-smokers, we found a significant (p< 0.0076, two-tailed Mann-Whitney U test) increase in the level of PE polymers in the smokers’ lungs as compared to healthy non-smokers. We also observed an increase in the levels of PC (p< 0.0947) and N6 (p < 0.0653) polymers within the lungs of healthy smokers, thus suggesting that smoking habits could affect the deposition/accumulation of select microplastic polymers within the lung tissues **(Figure 3A)**. Of note, we did not observe any significant variation in the accumulation of other microplastic polymers – PP, ABS, SBR, PMMA, PVC, PET, PU, PS, N66-in human lung tissues with the smoking status of the donor **(Supplementary Figure 2)**.

**Figure 3:**
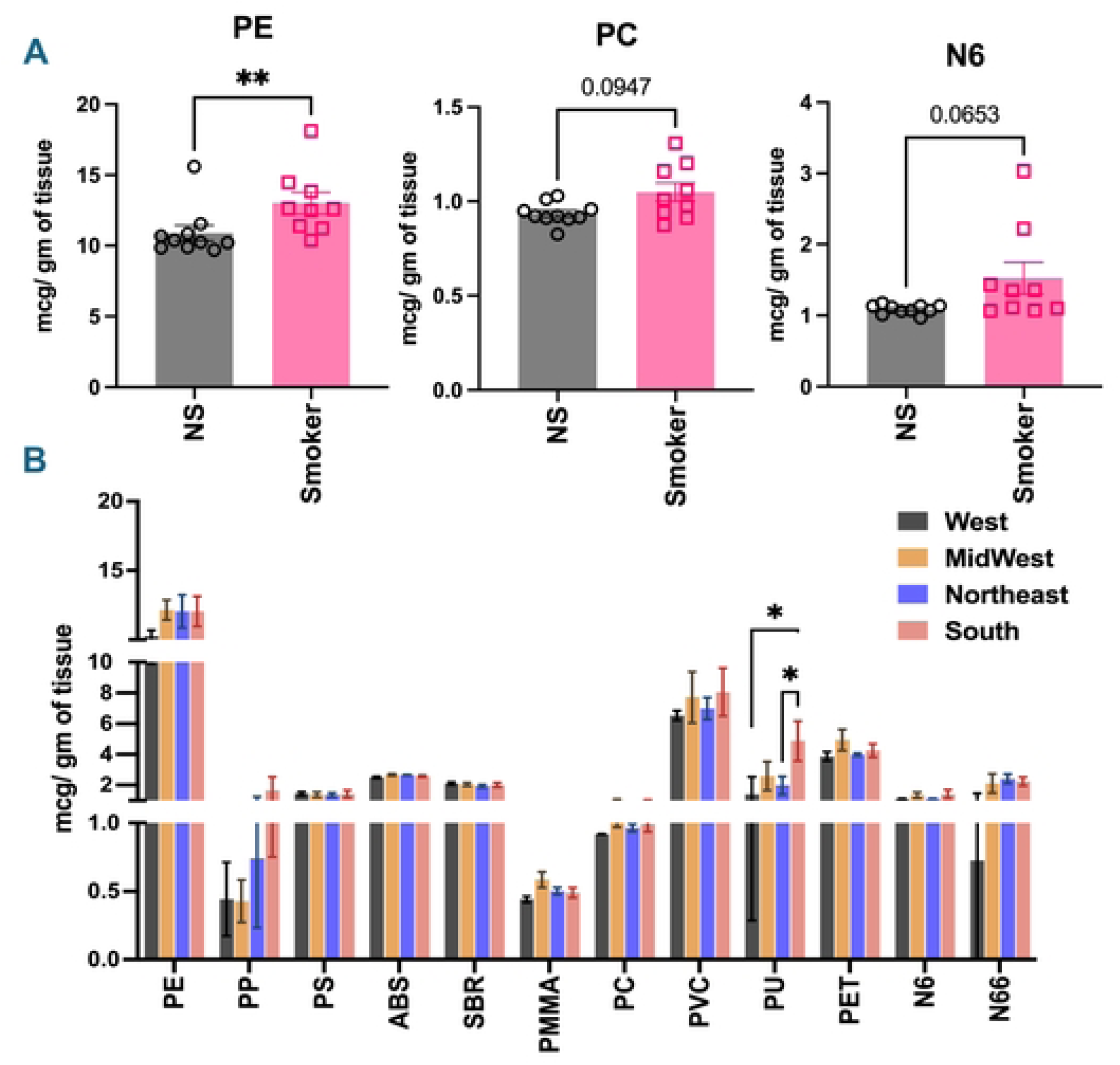
Significant increase in PE microplastics in the lung of smokers. Quantification of individual microplastics deposited in the human lung tissues was performed using Py-GC/MS. Bar graphs comparing the levels of PE, PC and N6 in the lungs of non-smokers (NS) and smokers. Data is represented as mean ± SEM for n = 9-10/group. SE: ** p< 0.01, per two-tailed Mann-Whitney U-test for pairwise comparisons **(A)**. Graph showing the distribution of individual microplastics per the region of residence of the donor, independent of the smoking history. Data is represented as mean ± SEM for n = 2-7/geographical region. SE: * p<0.05, per Two-way ANOVA with post hoc Tukey’s HSD test for multiple comparisons **(B)**.

A two-way ANOVA analysis was conducted to examine the effect of sex and smoking status on the accumulation of microplastics within the lungs. There was no statistically significant interaction between sex and smoking habit of donors on the distribution of 12 microplastic polymers being studied, as shown in **Table 2**. However, simple main effect analyses revealed that the levels of Poly (methyl methacrylate) (PMMA) accumulation were higher in males (p < 0.05) but showed no change with smoking status (p = 0.42). Similarly, we observed an increase in the accumulation of PE (p < 0.06), PC (p < 0.05) and PET (p < 0.05) in the smoker’s lung, with no significant variation with sex upon simple main effect analyses of smoking status amongst the male and female donor lungs.

**Table 2:**
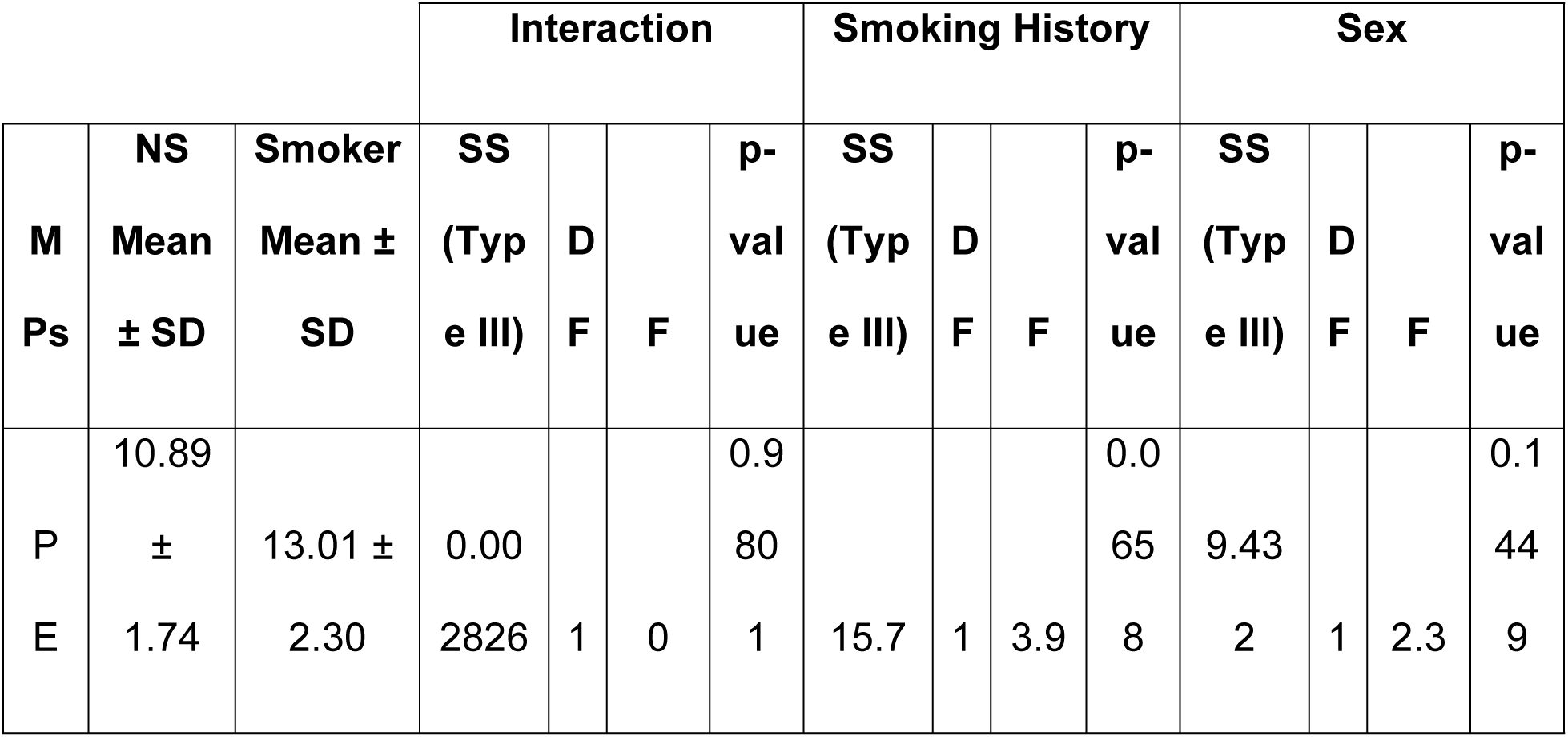
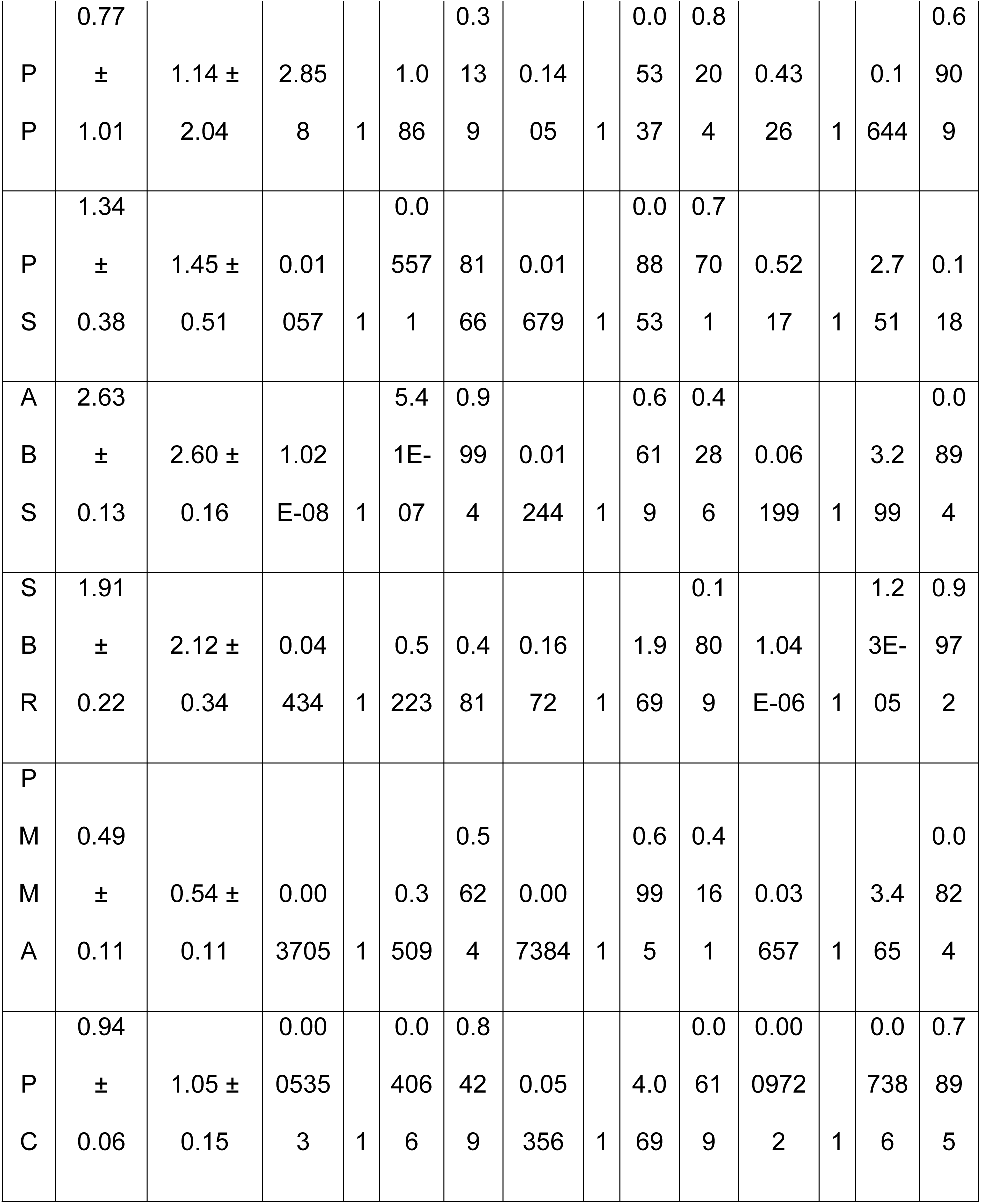
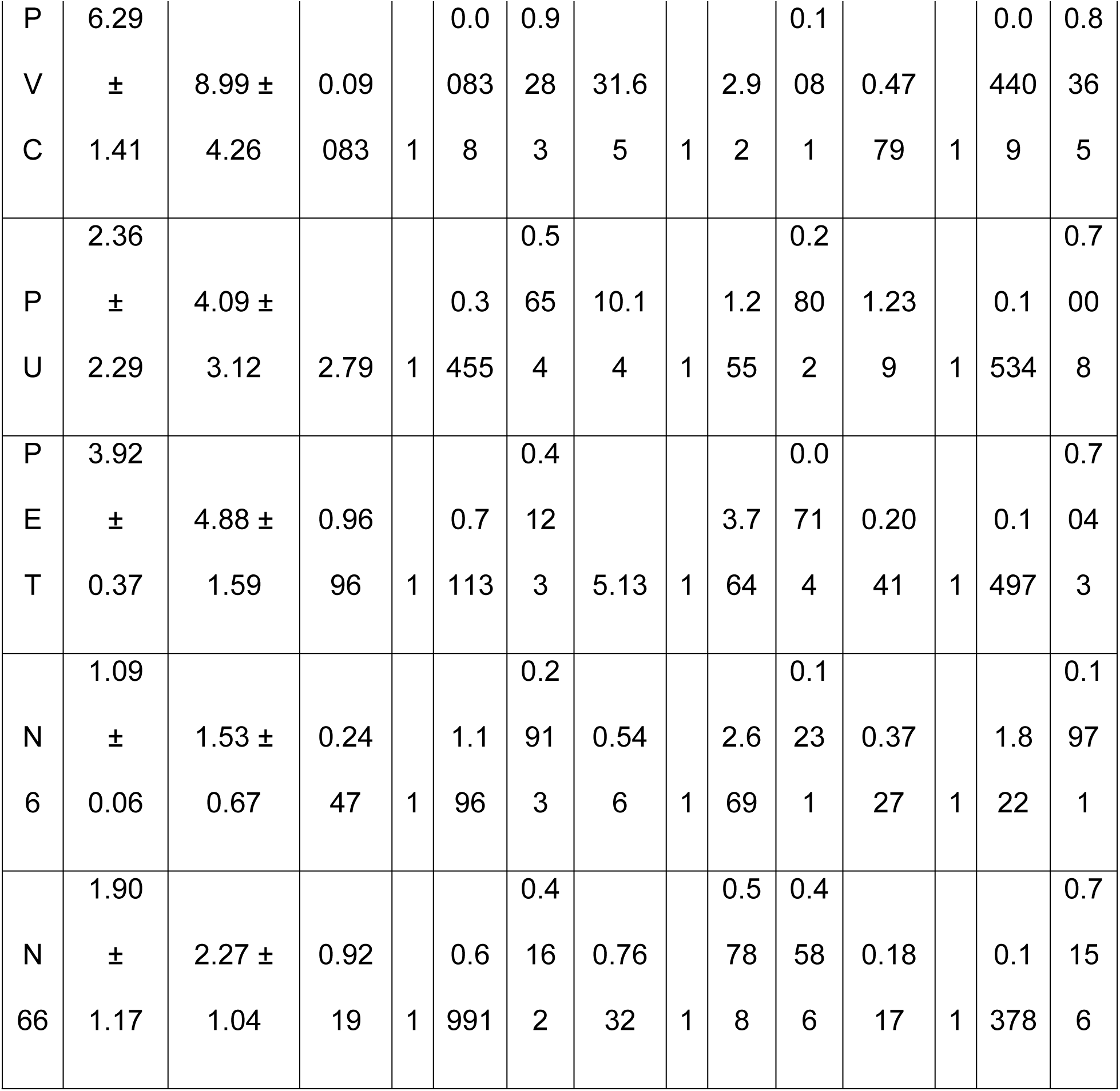
Table showing the average levels of microplastics (MPs; mcg per gm of lung tissue) detected in the lungs of non-smokers (NS) and smokers, and two-way statistics using smoking history and Sex as independent variables.

We next divided the donors (independent of smoking status) per the sampling sites into 4 regions within the United States. The four regions were: (a) Northeast: including Connecticut, Maine, Massachusetts, New Hampshire, New Jersey, New York, Pennsylvania, Rhode Island, and Vermont, (b) Midwest: including Illinois, Indiana, Iowa, Kansas, Michigan, Minnesota, Missouri, Nebraska, North Dakota, Ohio, South Dakota, and Wisconsin, (c) South: including Alabama, Arkansas, Delaware, District of Columbia, Florida, Georgia, Kentucky, Louisiana, Maryland, Mississippi, North Carolina, Oklahoma, South Carolina, Tennessee, Texas, Virginia, and West Virginia, and (d) West: including Alaska, Arizona, California, Colorado, Hawaii, Idaho, Montana, Nevada, New Mexico, Oregon, Utah, Washington, and Wyoming. Interestingly, we found a statistically significant increase in the levels of polyurethane (PU) identified in the lungs of donors from South as compared to Northeast (p < 0.024) and West (p < 0.0397) **(Figure 3B)**, thus suggesting that the polymer accumulation might also depend on the sampling site.

We also divided our donor lungs into three categories: < 30, 31-55, and >55 years, based on the age of each donor. One-way ANOVA analyses did not show any significant change in the levels of identified microplastic polymer with age in the lung tissues from human subjects included in this study **(Supplementary Figure 3)**.

### Sex-dependent increase in the levels of cadmium in smokers’ lungs

Considering reports suggest increased absorption of heavy metals upon accumulation of microplastics through environmental studies (Akhbarizadeh, Moore and Keshavarzi 2018, Abidli, Akkari et al. 2021, Liu, Shi et al. 2021), we hypothesized that similar effects could be expected within the mammalian system. To study this, we first quantified the levels of metals within the lungs of the same set of donors used for microplastic quantification using ICP-MS. Of the 53 elements analyzed through ICP-MS, 15 elements (S, Na, Mg, Al, Si, P, K, Ca, Ti, Mn, Fe, Cu, Zn, Rb, and Cd) exhibited levels above 1 ng/mg of tissue weight. Of these, only cadmium (Cd) showed a significant (p = <0.0001 per two-tailed Mann-Whitney U-test) increase in the lungs of smokers as compared to healthy non-smokers **(Supplementary Figure 4)**. Upon performing two-way ANOVA analyses on this dataset, we identified a significant interaction (p< 0.001) between sex and smoking habit of donors on levels of cadmium identified in the lung tissues **(Table 3)**. Simple primary effect analyses showed a significant increase in the level of deposited cadmium in the smokers’ lungs (p< 0.0001), which was more pronounced for female smokers as compared to their male counterparts (p = 0.0168) **(Figure 4A).**

**Figure 4:**
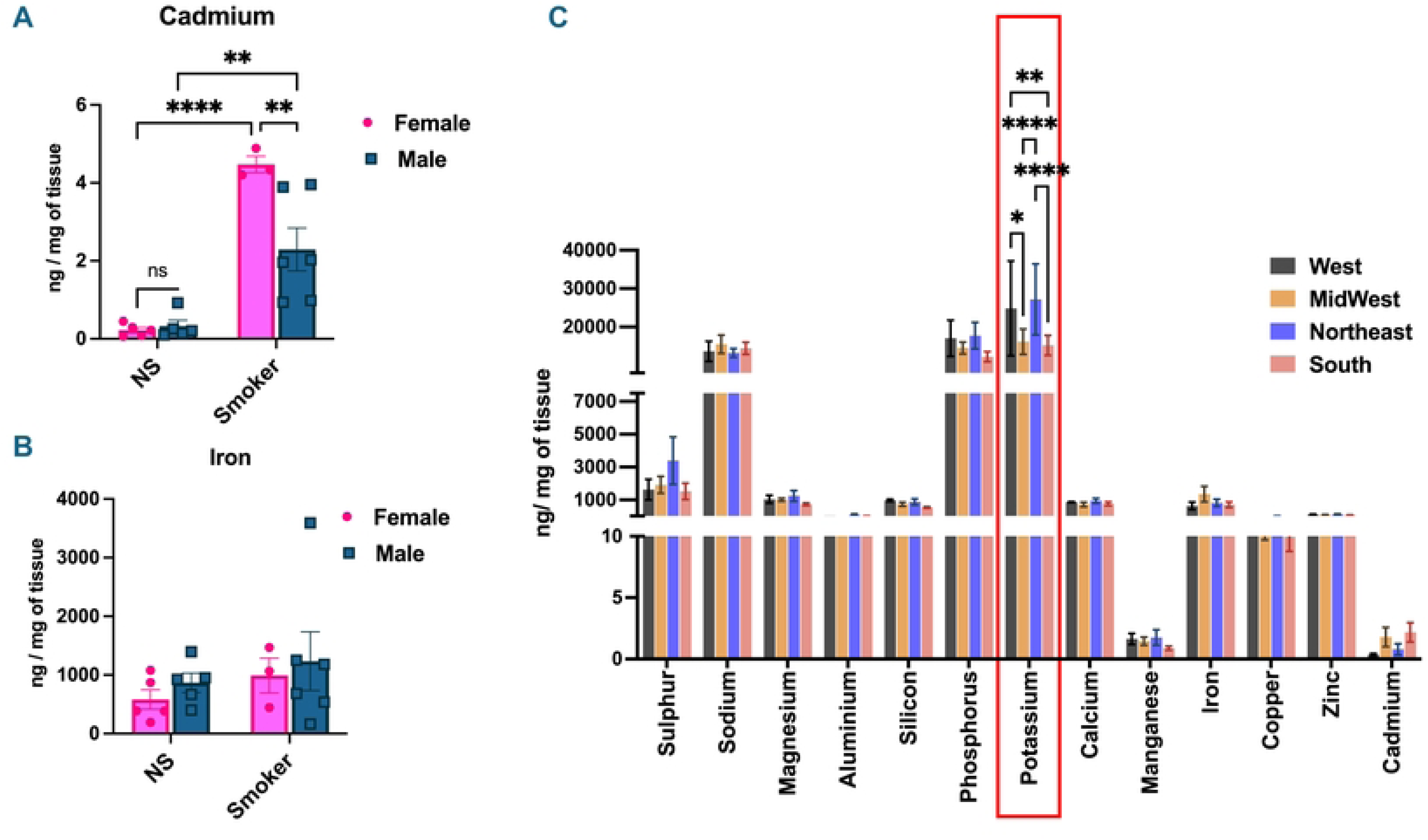
Sex-dependent increase in the levels of cadmium in smokers’ lungs. Levels of metals deposited in the lungs of healthy non-smokers (NS) and smokers were quantitated using ICP-MS. Bar graph showing levels of Cadmium **(A)** and Iron **(B)** in the lungs of healthy NS and smokers. Data is represented as mean ± SEM for n = 3-6/group. SE: ** p< 0.01 and **** p < 0.0001, per Two-way ANOVA with post hoc Tukey’s HSD test for multiple comparisons. Graph showing the distribution of select metals per the region of residence of the donor, independent of the smoking history. Data is represented as mean ± SEM for n = 2-7/geographical region. SE: ** p<0.01 and **** p< 0.0001, per Two-way ANOVA with post hoc Tukey’s HSD test for multiple comparisons **(C)**.

**Table 3:**
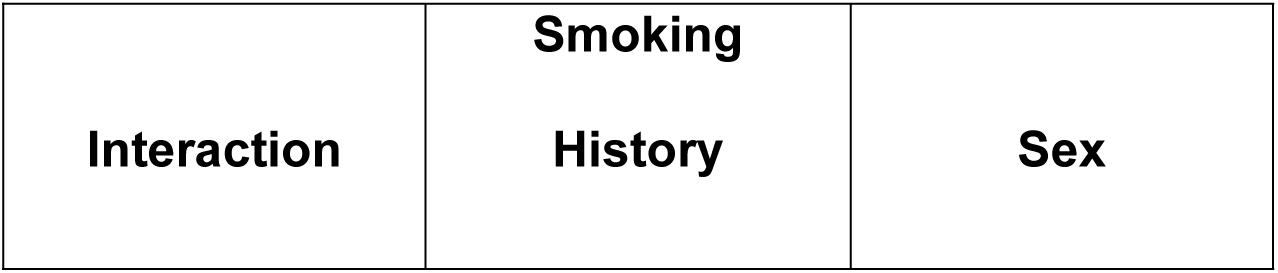
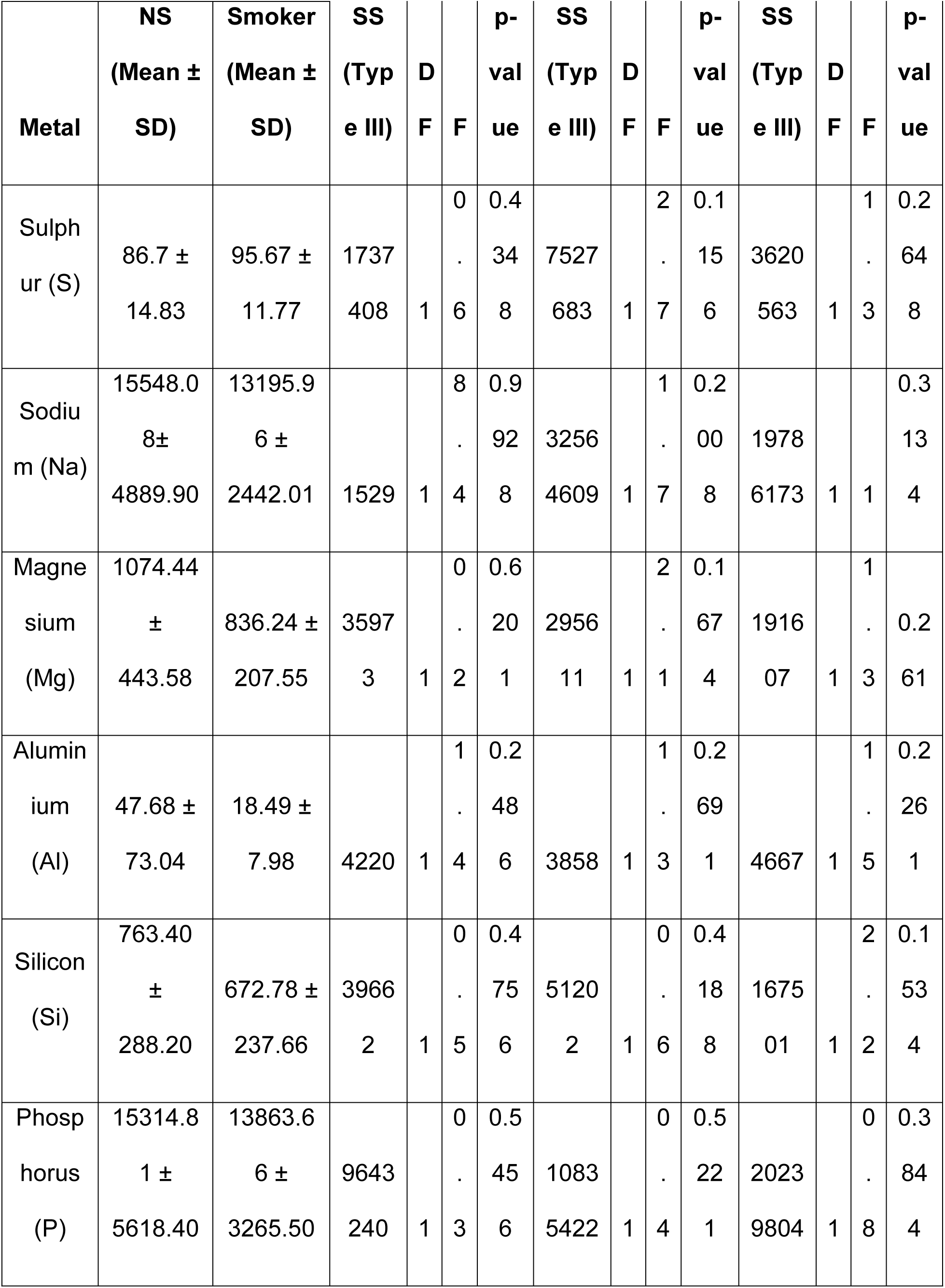
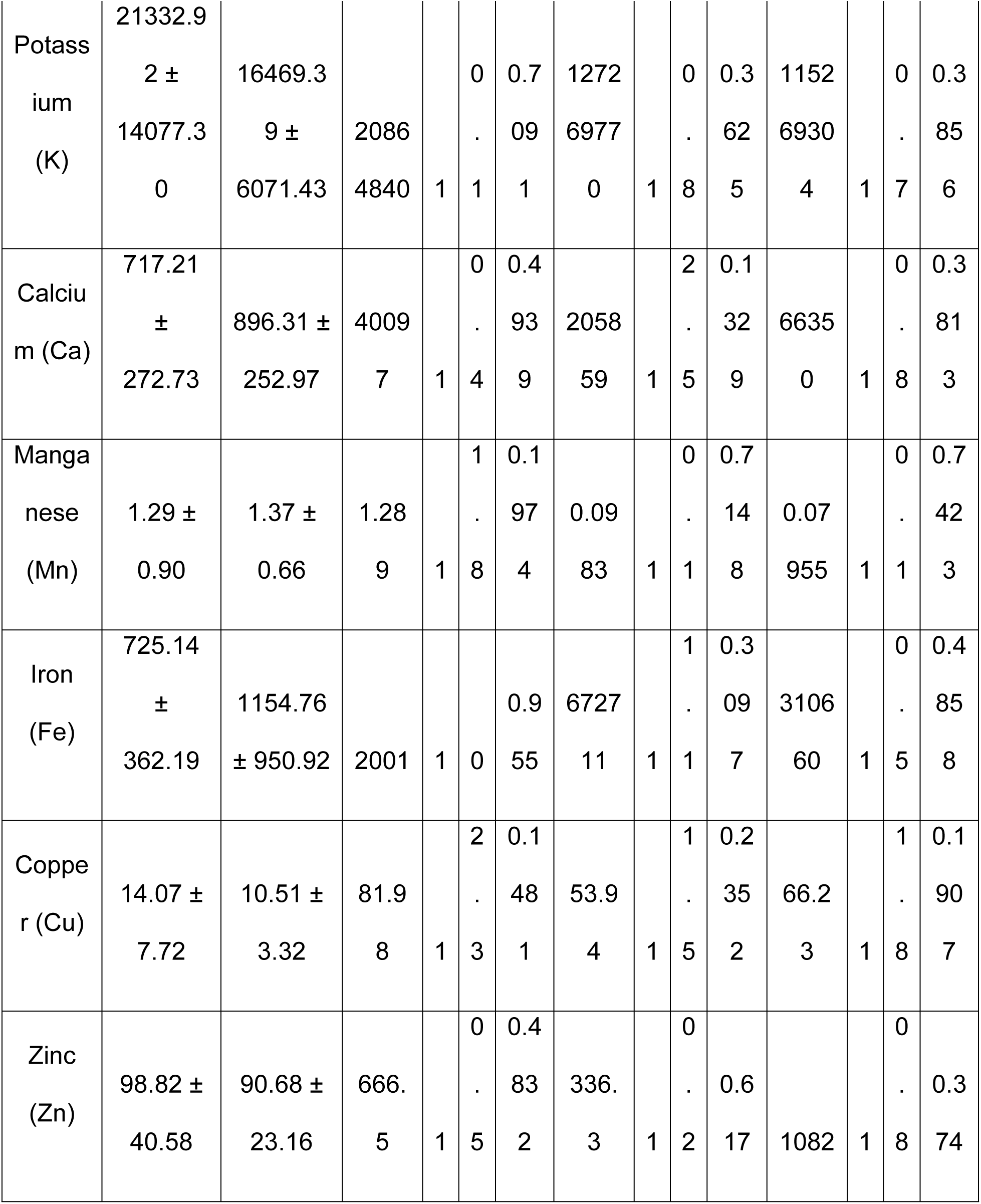
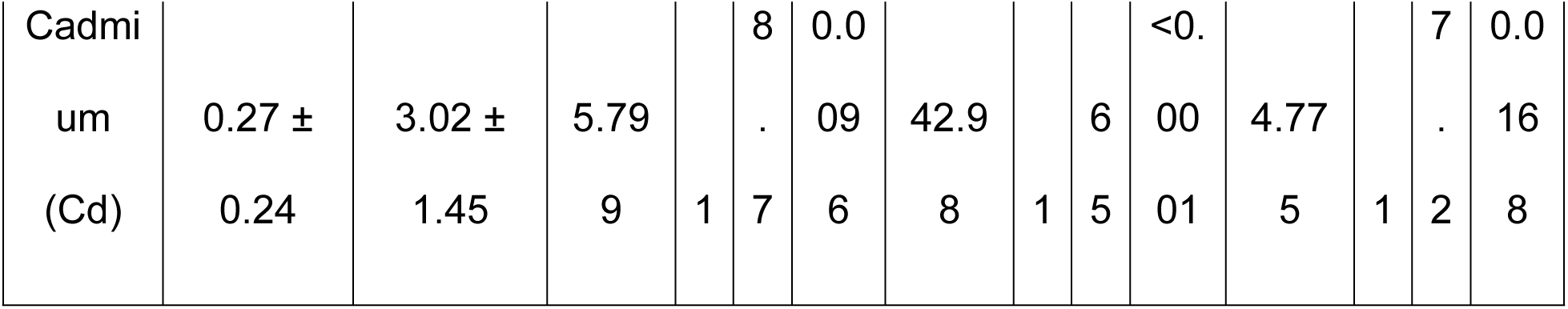
Table showing the average levels of metals (ng/mg of lung tissue) detected in the lungs of non-smokers (NS) and smokers, and two-way statistics using smoking history and Sex as independent variables.

Though we did not identify statistically significant interactions between sex and smoking habit on the deposition of any other heavy metals within the lungs in this study, sex-related trends still emerged that have been shown in **Supplementary Figure 5.** Further analyses of these changes with a larger sample size may reveal interesting patterns involving the accumulation of heavy metals within human lungs in a sex-dependent manner.

Upon analyzing the levels of metal deposited in the lungs based on the geographical location of the donors, we found significant variation in the levels of potassium (K) within the lungs based on their region of residence. Apart from K, the distribution of the other metals in the lungs remained unaffected by the geographical location of the donor in our study **(Figure 4C).**

### Levels of PE polymer accumulated in the lung tissue correlated significantly correlated with the cadmium levels

To establish whether the microplastic accumulation correlated with the heavy metal deposition within human lungs, we performed Pearson’s correlation of the levels of PE polymer and cadmium identified in the lungs of each donor. Interestingly, we observed a strong correlation (R^2^ = 0.2162, p < 0.05) between the amounts of accumulated PE and levels of cadmium in the lungs of human donors **(Figure 5A)**, thus proving that the presence of microplastics within the lung tissue might impact the absorption/accumulation patterns of various heavy metals that could have probable implications on the occurrence of chronic lung diseases in humans. As increased cadmium levels are known to affect the iron homeostasis in cells (Silver, Lozoff and Meeker 2013, Fujiwara, Lee et al. 2020, Guo, Zhang et al. 2024), we also studied if there exists any correlation between the levels of Cd and iron quantified in the donor lungs. Of note, simple main effect analyses of two-way ANOVA results found no significant change in the levels of Fe in the lungs of smokers as compared to non-smokers (p = 0.31). Similarly, the levels of Fe within the lungs were not found to vary with sex (p= 0.49) in our study **(Figure 4B).** However, though not significant, the levels of Cd showed moderate correlation (R2 = 0.2001, p< 0.05) with the levels of Fe identified within the lungs of human donors **(Figure 5B)**, thus supporting the claims in literature suggesting increased ferroptosis in relation to cadmium exposure in humans.

**Figure 5:**
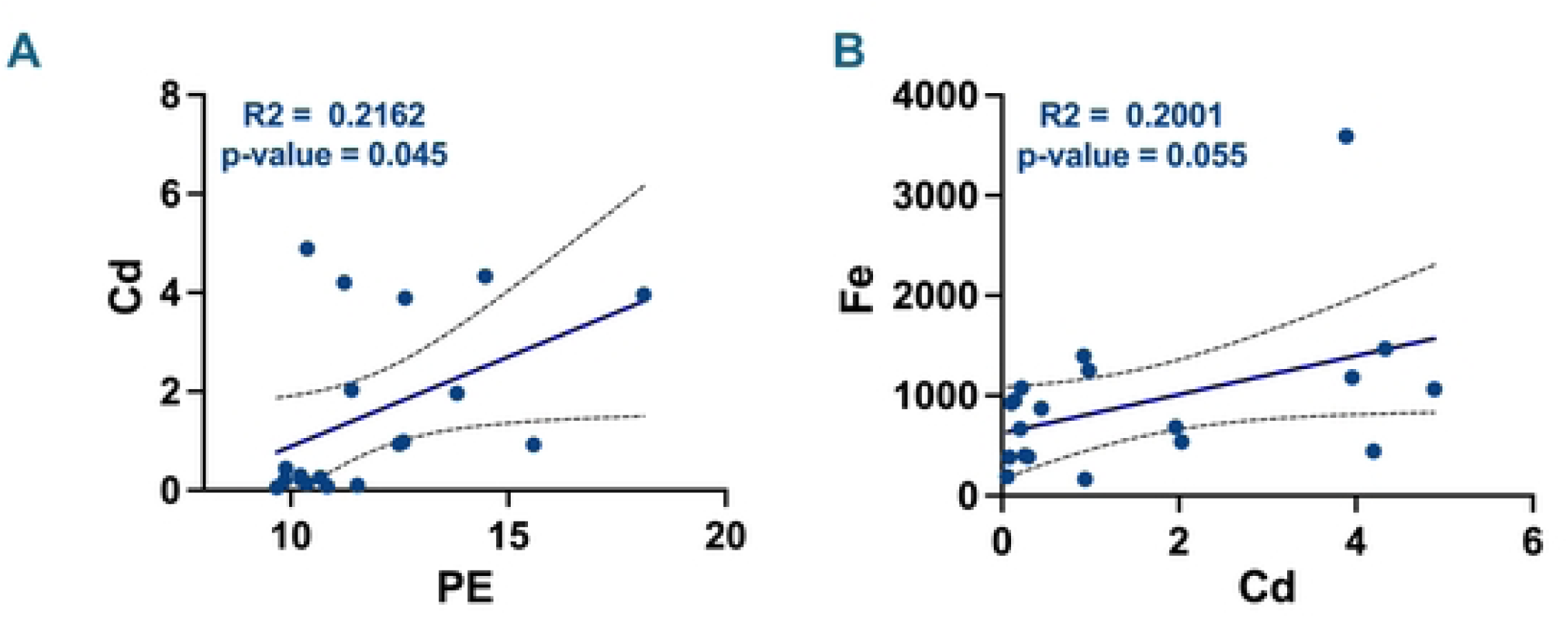
Levels of PE are significantly correlated with the Cadmium deposition in the lungs. Plots showing Pearson’s correlation between the levels of Cadmium (Cd) versus PE **(A)** and iron (Fe) versus cadmium (Cd) **(B)** in lungs from human donors.

## Discussion

The rapid increase in global usage of plastics has led to their omnipresence and the emergence of the pressing environmental issue of MNP pollution. The versatility, durability and low production cost of plastics make them an ideal choice for a wide array of applications in the twentieth century (Xu, Lin et al. 2022, Vasse and Melgert 2024). However, reports of the bioaccumulation of MNPs in body fluids (blood, sputum and BALF) (Huang, Huang et al. 2022, Lu, Li et al. 2023, Qiu, Lu et al. 2023, Wang, Lu et al. 2023), tissues (liver, colon, lung and brain), (Amato-Lourenço, Carvalho-Oliveira et al. 2021, Ibrahim, Tuan Anuar et al. 2021, Horvatits, Tamminga et al. 2022, Jenner, Rotchell et al. 2022, Massardo, Verzola et al. 2024, Nihart, Garcia et al. 2025) and placenta (Ragusa, Svelato et al. 2021, Garcia, Romero et al. 2024, Jochum, Garcia et al. 2025) from humans have emerged as a global concern with a pressing need for deeper scientific investigation and urgent public health scrutiny.

The airborne MNPs can originate from primary (wear and tear of synthetic textiles, abrasion of synthetic rubber tires, urban and household dust) or secondary (photoaging, wind erosion, and biodegradation of bulk plastics) sources. They usually differ in size (2.5-100 µm), shape (fibers and fragments) and chemical compositions (PP, PE, PS, and PET) (Zhao, Zhou et al. 2023, Yakovenko, Pérez-Serrano et al. 2025). Highly sensitive techniques like micro-Fourier Transform Infrared (µFTIR) spectroscopy and Raman spectroscopy have enabled the visualization and characterization of the microplastics in the lung tissues and lower respiratory tract (Jenner, Rotchell et al. 2022, Konings, Zada et al. 2024). However, there is a dearth of information regarding the accumulation patterns of microplastics in the lungs of smokers.

Recent data suggest that cigarette butts may release large volumes of microplastics with low rates of degradation. The traditional cigarettes are composed of plastics which could be a source of higher levels of MNPs in smokers’ lungs (Belzagui, Buscio et al. 2021, Shen, Li et al. 2021). Our study supports this viewpoint with proof of increased levels of total average microplastics in the lungs of smokers as compared to age and sex matched controls. Our findings corroborate the results from a 2023 study by Lu et al that showed higher concentrations of total microplastics, polyurethane, and silicone particles in the bronchoalveolar lavage fluid (BALF) samples from smokers (n=17) as compared to the control (n=15) samples collected in Zhuhai city, China (Lu, Li et al. 2023). We found increased levels of PE, PU and N6 in the lungs of smokers as compared to controls. Importantly, none of these polymers is a constituent of cigarette filters that are made up of cellulose acetate (Shen, Li et al. 2021). Thus, the increased accumulation of these microplastics could be due to the higher affinity of constituents of tobacco smoke towards these polymers. Many studies have reported the interaction of MNPs and heavy metals, which could be a probable cause of bioaccumulation of specific MNP polymers in smokers (Liu, Shi et al. 2021, Maity, Biswas et al. 2021). Multiple biophysiochemical and environmental factors, including size, surface charge, crystallinity, pH, and temperature, affect the adsorption of heavy metals to the surface of MNPs (Maity, Biswas et al. 2021). We report increased levels of cadmium accumulation in the lungs of smokers that correlated with the bioaccumulation of PE. While there is no evidence of the synergistic effects of MNPs and Cd in the context of lung function, Wen et al (2018) showed severe oxidative stress and innate immune responses in Amazon discus fish when exposed to a mixture of PS microplastic and Cd. Importantly, no such response was noted when the fish was treated with single toxicants (Wen, Jin et al. 2018). In a similar study aimed to understand the combined effect of exposure to Cd and microplastics on carp, the authors found significant changes in the plasma biochemical parameters and intrinsic immunological factors following 30-day exposure (Banaee, Soltanian et al. 2019). This evidence supports our hypothesis that the combined effects of deposition of MNPs and heavy metals could result in a rapid progression of chronic lung diseases such as COPD/ILD.

Previous studies have shown dose-dependent induction of oxidative stress, cellular senescence, apoptosis, and p38 phosphorylation-mediated NF-κB activation upon exposure of A549 cells to PS/PP microplastics (Woo, Seo et al. 2023, Milillo, Aruffo et al. 2024). Another study proved that exposure of murine macrophages to PS-microplastics leads to an increase in glycolytic respiration, but a reduction in mitochondrial respiration (Merkley, Moss et al. 2022). Furthermore, exposure of human macrophages to sulfate-modified nano plastics resulted in an enhanced accumulation of lipid droplets, leading to differentiation of macrophages to foam cells (Florance, Chandrasekaran et al. 2022). Taken together, these studies prove that different categories of microplastics have a varied cellular response. However, research in this area is still exploratory, and more work using relevant categories of microplastics and associated toxicants is important to understand the implications of inhalation of MNPs on lung pathologies.

While we show evidence of interaction between MNPs and heavy metals in the lungs of smokers, this could be one of the many chemical interactions leading to adverse toxic effects on lung health. For example, MNPs can interact with transition metals in lung epithelial lining fluid, leading to redox-dependent changes via Fenton’s/Haber-Weiss-like reactions. Polyamides have shown high affinity towards per- and polyfluoroalkyl substances (PFAS) found in nonstick cookware coatings, packaging, waterproof clothing, leather, and textile coatings, which have not been investigated in this study (Freilinger, Kappacher et al. 2025). Just like smoking, vaping could also lead to increased deposition of microplastics with heavy metals (due to heating of the coil) into the lungs of regular e-cig users (Wagner, Chen and Vrdoljak 2020).

Py-GC/MS has been implemented in recent studies (10, 13), ostensibly to assess the concentration of polymer solids in human tissues. However, this is an indirect method that entails meaningful uncertainty. The quantities determined for each polymer are based on representative pyrolyzate ions that are typical of pristine polymers, but not necessarily optimal for aged, environmental polymers. Furthermore, extraction of plastics from a complex biological matrix remains an important area of research with no consensus method at present. The addition of the cyclohexane wash after ultracentrifugation was included in this study to more aggressively remove lipids, but it may also reduce the organic corona of aged MPs. The field is evolving rapidly, and continued innovation is needed to improve confidence.

It will be interesting to characterize the size, shape, and polymer type deposited in the distal and peripheral airways. Given the evidence of bioaccumulation of MNPs with heavy metals in the lungs of individuals with a smoking history, future studies will need to investigate the nature of their interaction and its impact on immune responses and epithelial responses in the lungs. Overall, our study proved for the first time that smoking results in an increased accumulation of PE-microplastics within the human lungs that correlated with the levels of Cd, which may have implications for the progression of chronic lung ailments amongst smokers.

## Acknowledgements

We would like to extend our gratitude to the families and friends of the lung donors who agreed to provide human tissues for research.

We thank Dr. Matthew Rand and Thomas Scrimale at the Elemental Core Analyses facility at URMC for metal quantification.

## Data Availability

Source data are provided with this paper.

## Author’s contributions

GK, MIF and FE planned and conducted the experiments; AD and MG quantified the levels of MNPs in the isolated samples; GK analyzed the results, wrote and edited the manuscript; All authors reviewed the data and manuscript, and IR conceived and procured funding for the project.

